# Healthy human induced pluripotent stem cell-derived cardiomyocytes exhibit sex dimorphism even without the addition of hormones

**DOI:** 10.1101/2024.05.29.596547

**Authors:** Sophie E. Givens, Abygail A. Andebrhan, Eric G. Schmuck, Aimee Renaud, Juan E. Abrahante, Noah Stanis, James R. Dutton, Brenda M. Ogle

**Affiliations:** Biomedical Engineering, University of Minnesota, Minneapolis, MN, USA; Stem Cell & Regenerative Medicine Center, University of Wisconsin-Madison. Madison, WI, USA; Stem Cell Institute, University of Minnesota, Minneapolis, MN, USA; Informatics Institute, University of Minnesota, Minneapolis, MN, USA; Department of Pediatrics, University of Minnesota, Minneapolis, MN, USA; Institute of Engineering in Medicine, University of Minnesota, MN, USA; Department of Genetics, Cell Biology and Development, University of Minnesota, Minneapolis, MN, USA

## Abstract

Human induced pluripotent stem cell-derived cardiomyocytes (hiPSC-CM) are a valuable cell type for studying human cardiac health and disease *in vitro*. However, it is not known whether hiPSC-CM display sex dimorphism and therefore whether sex should be incorporated as a biological variable in *in vitro* studies that include this cell type. To date, the vast majority of studies that utilize hiPSC-CM do not include both male and female sex nor stratify results based on sex because it is challenging to amass such a cohort of cells. Here we generated three female and three male hiPSC-lines from adult left ventricular cardiac fibroblasts as a resource for studying sex differences in *in vitro* cardiac models. We used this resource to generate hiPSC-CM and maintained them in basal media without exogenous hormones. Functional assessment of CM showed enhanced calcium handling in female-derived hiPSC-CM relative to male. Bulk RNA sequencing revealed over 300 differentially expressed genes (DEG) between male and female hiPSC-CM. Some of the DEG are X and Y-linked genes and many are implicated in cardiac health and disease including potassium channels which could account for net differences in calcium handling shown here. Gene ontology analysis of DEG showed distinct differences in pathways related to cardiac pathology including cell-cell adhesion, metabolic processes, and response to ischemic stress. These findings highlight the importance of considering sex as a variable when conducting studies to evaluate aspects of human cardiac health and disease related to cardiomyocyte function.

## Introduction

Cardiovascular disease (CVD) is the leading cause of mortality and morbidity globally^1^ but historically it has been challenging to study *human* CVD at the cellular level. Over the past decade stem cells, particularly human induced pluripotent stem cells (hiPSC), have become a source for creating *in vitro* model systems to study *human* CVD. Robust protocols for yielding heart muscle cells or cardiomyocytes have propelled this approach^2–4^. Using small molecule WNT modulation, many hiPSC-CMs can be generated in a dish^2^ and purified using metabolic selection^5^. These cells represent a versatile, ethical, and patient-specific model system for studying CVD. However, the use of this model system rarely considers sex as a biological variable though sex is a key mediator of CVD onset, progression, and prognosis in the clinic.

When developing models of CVD, consideration of sex dimorphism is essential as clinical and animal studies have revealed stark sex differences in CVD onset and progression^6–9^. For example, women with ischemic heart disease (IHD) often have microvasculature and endothelial dysfunction and are more prone to spontaneous coronary artery dissection^10,11^. In contrast, men experience higher rates of coronary artery dissections linked to IHD and are more susceptible to ischemia-reperfusion injury^10,12,13^. Additionally, studies have shown distinct differences in myocardial response to pressure overload, a common precursor of heart failure, in female and male animals^14–16^. Rat studies have shown that females typically exhibit diastolic dysfunction characterized by decreased compliance of the heart muscle, for which no mechanistic therapies have been established, and males more often develop systolic dysfunction with decreased ejection fraction and increased fibrosis, for which therapies have been established^17^. Heart failure itself also shows a sex-dependent divergence; HF with preserved ejection fraction, coupled with increased ventricular stiffness, is more common in women, whereas men are more afflicted by HF with reduced ejection fraction, leading to ventricular dilation^6^. Arrhythmias demonstrate similar sex-specific occurrences. Women have higher instances of acquired long-QT syndrome and Torsades de Pointes, while men are at increased risk for atrial fibrillation and sudden cardiac arrest^7^. Most sex differences detected in CVD thus far have been attributed to genetic, epigenetic, or hormonal factors^8,9^. Since these factors can be hard to uncouple, especially in a human system and at the cellular level, there is still much to learn about sex dimorphism in human CVD. Given limited access to human cells and tissues, hiPSCs are an enabling tool to study human sex dimorphism in the cardiovascular system.

Numerous models of CVD have been generated using hiPSC-CM since their first derivation more than a decade ago^18,19^. These include disease models of IHD and ischemia reperfusion injury^20–22^ and even more commonly studies of cardiac arrythmias are done using hiPSC-CM; these include hiPSC-CM models of long-QT, Torsades de Pointes and atrial fibrillation which have all been seen to be sex dimorphic in the clinic^23–27^. Interestingly, there has been very little consideration of sex dimorphism in these models. There have only been a few studies utilizing stem cell-derived CM that look at sex differences. Among them, one study looked at sex differences in hESC-CM hypertrophy in response to isoproterenol using one female and one male line. Results showed that female CM undergo hypertrophy more slowly in response to isoproterenol indicating cellular level differences in CM hypertrophy without the addition of hormones^28^. In another study, three female and three male hiPSC-CM lines from different ethnic origins were tested for sensitivity to various pharmacological agents^29^. This study found that hiPSC-CM from females were more sensitive to inward rectifying potassium channel (I_Kr_) blockers than hiPSC-CM from males. This finding was confirmed in another study using hiPSC-CM to study drug-induced Torsades de Pointes; in this study female hiPSC-CM exhibited an increase in long-QT generation when exposed to I_Kr_ blockers^30^. The exogenous addition of hormones affected hiPSC-CM electrophysiology but did not change the sensitivity of the female hiPSC-CM to I_Kr_ blockers. This was attributed to increased expression of KCNE1, a critical cardiac potassium channel, in male hiPSC-CM. These studies were significant as they showed that the potential mechanism for sex dimorphism in drug-acquired Torsades de Pointes is not hormone-mediated. These studies support the possibility that other cellular differences in human CM are present without hormones which are commonly thought to be the major mediator of sex dimorphism in CVD. The few studies that look at sex differences in hiPSC-CM model systems motivate future work wherein hiPSC-CM are used to determine the mechanism of CVD sex dimorphism in a controlled manner.

Sex differences have also been reported in heart development, as well as heart function in healthy controls. A recent study examining cell composition in various regions of the heart found that females tend to have a higher percentage of ventricular cardiomyocytes and this is in turn associated with a lower proportion of fibroblasts (FBs)^31^. This indicates differences in cellular composition in the healthy male and female heart. Functionally, rat studies indicate that females exhibit a smaller ejection fraction and reduced fractional shortening relative to their male counterparts^32,33^. Further, animal studies show increased calcium-handling in cardiomyocytes of the female heart^34,35^. Additionally, there are significant transcriptional differences between male and female rat ventricular cardiomyocytes^32^. These data were recently bolstered by human single cell sequencing data at different developmental time-points. It showed that female and male cardiomyocytes, had the largest number of differentially expressed genes at fetal, young, and adult stages when compared to all the other cell types in the heart^36^. The high prevalence of sex differences in both cardiac health and disease provides a strong rationale for the consideration of sex as a biological variable in all studies wherein cardiomyocyte health or dysfunction is under evaluation.

Here we developed a cohort of male and female hiPSC lines from six healthy donors as a resource to enable the study of sex dimorphism with cardiac health and disease. The hiPSCs were made from the same cell source, at a similar donor age, and with the same reprogramming methodology to control for epigenetic noise^37,38^. Though other larger hiPSC resources exist many still cobble together lines from multiple cell sources that were derived in various ways by multiple labs. Additionally, the lines were made from cardiac fibroblast as they can yield more functionally relevant hiPSC-CM^39,40^. To better understand if there are sex differences in hiPSC-CM without the exogenous addition of hormones, we differentiated hiPSC-CM from all of the hiPSC-lines generated and maintained them in basal CM media (RPMI/B27 with insulin). We opted to conduct our comparison without the addition of sex hormones to mimic the setting of most basic bench research utilizing hiPSC-CM and to uncouple hormone regulation from basal function of hiPSC-CM.

## Results

### Sex is underreported in studies that utilize hiPSC-CM

To determine whether sex has been considered in hiPSC-CM research, we looked at literature over the past 13 years, starting in 2010 when hiPSC-CM differentiation protocols first began to appear. We searched for all papers pertaining to hiPSC-CMs, with the query “(Human Induced Pluripotent Stem Cell derived Cardiomyocytes) OR hiPSC-CMs OR (human iPSC-CMs).” As expected, the number of papers drastically increased from 2010 to 2023 (**Figure 1A**). This search was refined for any papers mentioning hiPSC-CM and sex, male, or female to determine if one of these terms was even mentioned in the manuscripts. Only around 20 percent of the papers mentioning hiPSC-CM also mentioned the terms “sex,” “male” or “female” (**Figure 1B**). Further narrowing our focus, we determined that only 1% of papers about hiPSC-CM mentioned “sex,” “male,” and “female” at the same time (**Figure 1C**), indicating it is likely that less than 1% of the hiPSC-CM literature uses both male and female lines or stratifies results by sex. To validate this, the top 50 hiPSC-CM primary research papers between the year of 2010 and 2023, if there were 50 papers for that year, on PubMed were analyzed more closely (**Figure 1D**). The papers were probed carefully to determine if 1) the sex of the hiPSC-line or lines used in the study were reported and 2) if the results were analyzed by sex. We observed a gradual increase in the percentage of papers reporting the sex of cell lines over this period (**Figure 1E**). Despite this increase, by 2022 only 70% of the papers reported the sex of the lines they used or reported the line origin in which the sex could be easily determined. Additionally, only three papers in this search analyzed their results by sex (**Figure 1F**). This suggests that studies rarely compared their findings between hiPSC-CM from both male and female donors. In summary, more papers are reporting the sex of hiPSC-CM utilized, but sex-related differences in these studies are not being evaluated.

**Figure 1.**
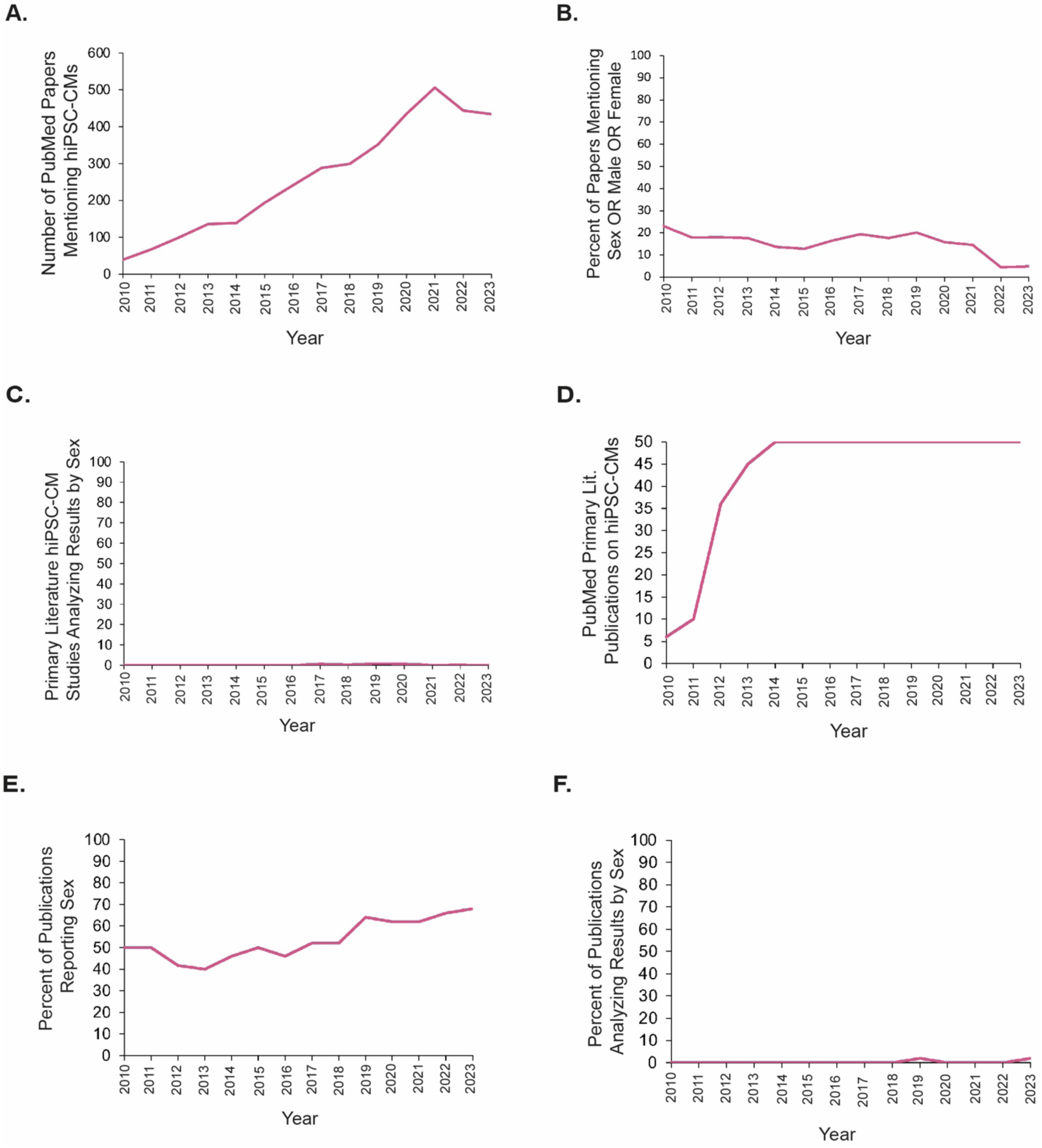
Sex is underreported and understudied in hiPSC-CM literature. A) A line graph illustrating the annual number of papers retrieved when searching “(Human Induced Pluripotent Stem Cell derived Cardiomyocytes) OR hiPSC-CMs OR (human iPSC-CMs)” on PubMed for the years 2010 to 2023. B) A line graph illustrating the annual number of papers retrieved when searching “((Human Induced Pluripotent Stem Cell derived Cardiomyocytes) OR hiPSC-CM OR (human iPSC-CMs)) AND (Sex OR Male OR Female)” in PubMed for the years 2010 to 2023. C) A line graph illustrating the annual number of papers retrieved when searching “((Human Induced Pluripotent Stem Cell derived Cardiomyocytes) OR hiPSC-CMs OR (human iPSC-CMs)) AND (Sex AND Male AND Female)” on PubMed for the years 2010 to 2023. D) A line graph displaying the number of papers more thoroughly assessed in (E) and (F) when searching “(Human Induced Pluripotent Stem Cell derived Cardiomyocytes) OR hiPSC-CMs OR (human iPSC-CMs)” on PubMed for the years 2010 to 2023. E) Represents the number of primary literature papers that reported the sex of the hiPSC-line and F) displays the number of primary literature papers that analyzed their results by sex when searching the query above on PubMed.

### Creating a cohort of male and female hiPSCs with consistent cell source and reprogramming method

To enable studies of human CM that allow for the stratification of results according to sex, human adult left ventricular fibroblasts from three female and three male lines were reprogrammed into hiPSCs using the Sendai virus (SeV) reprograming method (**Figure 2A**). Young adult donors between the ages of 21 and 34 years old with no known cardiac abnormalities were used (**Table 1**). Three lines were generated from individual colonies chosen from each donor and all 18 lines were confirmed to be SeV depleted via q-RT-PCR (**Table 1 and S1**). One line from each donor was assessed for pluripotency using q-RT-PCR for the pluripotency genes OCT4, SOX2, and NANOG in comparison to a previously established hiPSC-line, the CCND2-hiPSC line, as a positive control (**Figure 2B**)^41^. Representative brightfield images of line 1 (L1) from all 6 donors show the distinct colony morphology characteristic of hiPSCs (**Figure 2C**). The gene expression of the pluripotency marker OCT4 was validated at the protein level using flow cytometry showing > 95% Oct4+ cells in L1 from all six donors (**Figure 2D**). The cell surface pluripotency marker SSEA4 was determined to be present in all 6 cell lines as well (**Figure S1A**). To visually inspect the presence of the pluripotency markers in the F6, F7, M4, and M7 lines, co-expression of transcription factor Sox2 and the cell surface marker TRA-1-60 were detected by immunohistochemistry (**Figure 2E and Figure S2B**). Similarly, co-expression of the transcription factor Oct4 was detected together with the cell surface marker SSEA4 (**Figure S2C**).

**Figure 2.**
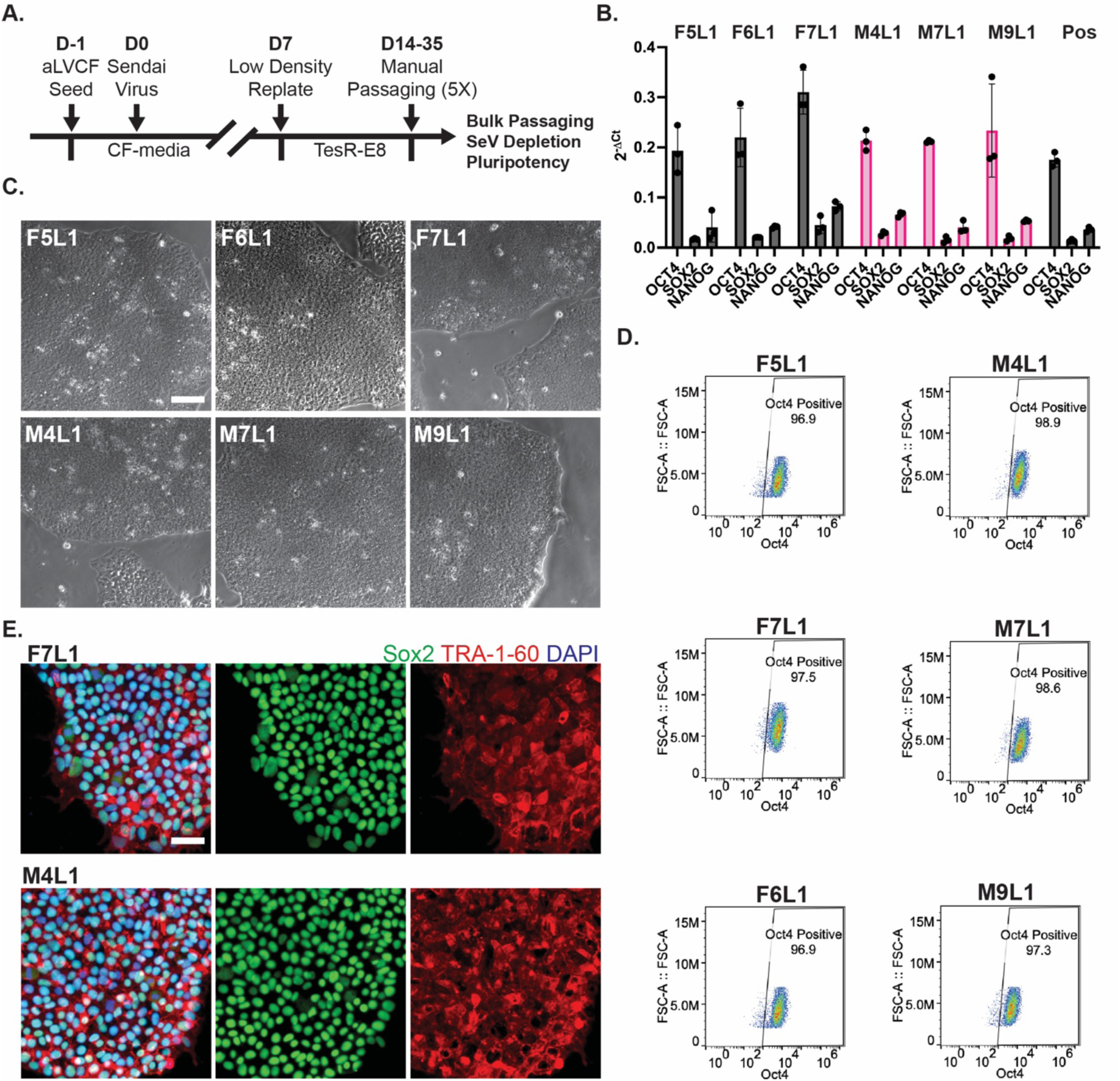
Adult left ventricular cardiac fibroblasts reprogramming into hiPSCs. A) Schematic depicting the Cyto-Tune 2.0 Sendai virus hiPSC reprogramming protocol used in this study. B) Pluripotency validation via q-RT-PCR of OCT4, SOX2, and NANOG as compared to a previously established line as a positive control (Pos). C) Brightfield images showing the characteristic hiPSC colony morphology in the adult left ventricular derived-hiPSCs from all 6 donors. D) Flow cytometry of the pluripotency marker Oct4 for all 6 donors with the gate decided based on an isotype-control. E) Representative IHC images of the pluripotency markers Sox2 and TRA-1-60 from one female (F7L1) and one male (M4L1) lines. Scale 50μm. For (B) the bar and error bars represent the mean ± STDEV and each dot represents one technical replicate for one passage of each hiPSC-lines after Sendai virus depletion where a line is defined as Sendai virus depleted when there is no amplification of q-RT-PCR Sendai virus primers for two passages.

**Table 1.** Adult left ventricular fibroblasts line donor age, line SeV depletion and karyotype.

| Line Name | Sex (F/M) | Age (Years) | Karyotype | SeV Depletion (Passage #) |
| --- | --- | --- | --- | --- |
| F5L1 | F | 26 | Normal XX (QC) | 18 |
| F6L1 | F | 34 | Normal XX (QC) | 16 |
| F7L1 | F | 36 | Normal XX (Full) | 12 |
| M4L1 | M | 21 | Normal XX (Full) | 12 |
| M7L1 | M | 21 | Normal XX (QC) | 8 |
| M9L10 | M | 27 | Normal XX (QC) | 8 |

To ensure that no chromosomal abnormalities were acquired during SeV reprogramming karyotype analysis was performed for 20 metaphases of one male (M4L1) and one female (F7L1) line (**Figure S2**). This is extremely important for hiPSC-lines being used to study sex differences as Y-linked, X-linked, and genes that escape X inactivation can be mechanistic mediators of sex dimorphism. Thus, a normal XY and XX karyotype is important for studying the dominant sex chromosome orientations. Quality control karyotype analysis examining 7 cells per line with the readout of “normal” or “abnormal” was conducted for L1 of the remaining donors. Karyotyping revealed a normal “XX” karyotype for all three female lines and a normal

“XY” karyotype for all three male lines (**Table 1 and S1**). Next, L1 from all 6 donors were differentiated into hiPSC-CM to evaluate genetic and functional characteristics.

### Female hiPSC-CM exhibit enhanced calcium transient properties

The female and male hiPSC-lines were differentiated into CM using small molecule WNT modulation and purified using lactate metabolic selection media (**Figure 3A**)^2,5^. On day 40, CM purity was assessed via flow cytometry for the CM-specific marker cardiac troponin T (cTnT) (**Figure 3B**). Cardiomyocytes displayed high purity, more than 95% cTnT positive, for all 6 cell lines (**Figure 3C**). The hiPSC-CM purity was not significantly different when analyzed by sex (**Figure 3D**).

**Figure 3.**
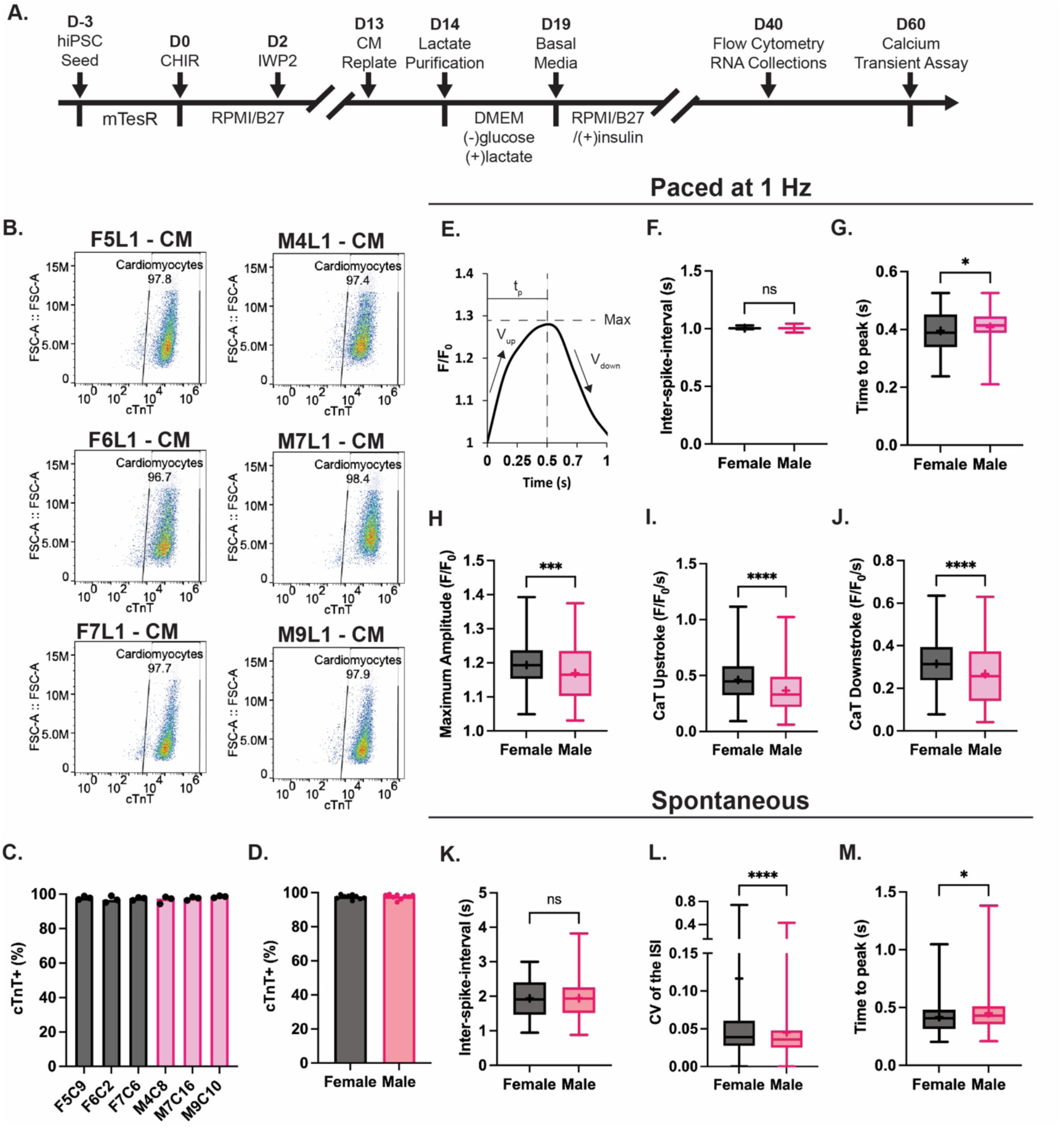
Female hiPSC-CM have enhanced calcium handling properties. A) Schematic depicting the hiPSC cardiomyocyte differentiation, purification, and the time-points of CM purity assessment, RNA collection, and calcium handling data acquisition. B) Representative flow cytometry data showing the cells on forward scatter (FSC) versus cardiomyocyte marker, cTnT on the x-axis. C) Quantification of the percent cardiomyocytes (cTnT+) by line and D) by sex where the bar represents the average between n = 3 independent experiments across all 6 lines. Each dot represents the given cTnT+ (%) for one replicate. E) Representative paced calcium handling trace (from M4L1-CM) showing some of the parameters assessed such as the time to peak (t_p_), the maximum amplitude (Max), and the maximum upstroke (V_up_) and downstroke (V_down_) velocities. Calcium transient of the male and female hiPSC-CM showing the F) inter-spike-interval, G) time to peak, H) maximum amplitude, and the calcium transient (CaT) I) upstroke and J) downstroke. Box and whisker plots of the spontaneous CaT parameters, the K) ISI L) the coefficient of variance (CV) of the ISI, showing the regularity of hiPSC-CM spontaneous CaT, and the M) time to peak, for the male and female hiPSC-CM. For (F-M) the box and whisker plots represent the median, upper, and lower quartiles while the (+) represents the mean. (F-M) The data is the average of 9 videos across three technical replicates for three independent experiments from all three female and all three male hiPSCs. *p < 0.05, **p < 0.01, ***p < 0.001, and ****p < 0.0001.

On day 60, calcium functional analysis was performed using Rhod-2AM dye. The calcium transient (CaT) inter-spike-interval, time to peak, maximum amplitude, downstroke, and upstroke velocities for each line were assessed (**Figure 3E**). CM were analyzed under both paced and spontaneous conditions. For the paced conditions, the hiPSC-CM were electrically stimulated at a frequency of 1Hz resulting in an inter-spike-interval (ISI) of 1s for both the female and male hiPSC-CM (**Figure 3F**). Controlling for beat rate allows for the accurate assessment of CM functionality via CaT time-to-peak and maximum amplitude as well as the CaT upstroke and downstroke velocities. The CaT time-to-peak is slightly but significantly decreased in the female hiPSC-CMs (**Figure 3F**) indicating a faster net release of calcium from the sarcoplasmic reticulum of female hiPSC-CM. Additionally, the CaT maximum amplitude as well as the CaT downstroke and upstroke velocities were significantly higher in female CMs (**Figure 3G-J**). Though there is a net increase in the female hiPSC-CM calcium handling when all of the lines are averaged it is important to note that there is substantial donor-to-donor variability (**Figure S4A-E**). The ANOVA post-hoc Tukey HSD comparison of means shows significant CaT line-to-line variability between all of the male and female donors (**Tables S4-S7**). The spontaneous calcium transient activity was assessed to determine if there were any intrinsic differences in CM beat rate or the regularity of beating as assessed by the coefficient of variance (CV) of the ISI. No significant difference in the spontaneous ISI were observed between the female and male hiPSC-CM (**Figure 3J**). However, the CV of the ISI was significantly greater in the female lines, indicating that the beat rate is more irregular in female hiPSC-CM compared to males (**Figure 3L**). Additionally, the decrease in time-to-peak in the female hiPSC-CM was seen in the spontaneous case as well (**Figure 3M**). The spontaneous calcium data also showed significant line-to-line variability (**Figure S4F-H**). The ANOVA post-hoc Tukey HSD comparison of means confirmed the line-to-line variability as well (**Table S8-S9**). The calcium functional analysis demonstrates that there are net functional differences between CMs derived from male and female hiPSC-lines. These results show a net increase in the calcium-handling ability of female hiPSC-CM in combination with a tendency for more irregular spontaneous beating frequencies.

### Bulk RNA sequencing of hiPSC-CM reveals distinct hierarchical clustering according to sex

On the same day that CM purity was validated, RNA was collected from all the male and female hiPSC-CM for bulk RNA sequencing. Three independent experiments from each hiPSC-line were analyzed. A heat map generated from the expression profiles of all 18 samples displayed a distinct hierarchical clustering between male (M) and female (F) hiPSC-CMs (**Figure 4A**).

**Figure 4.**
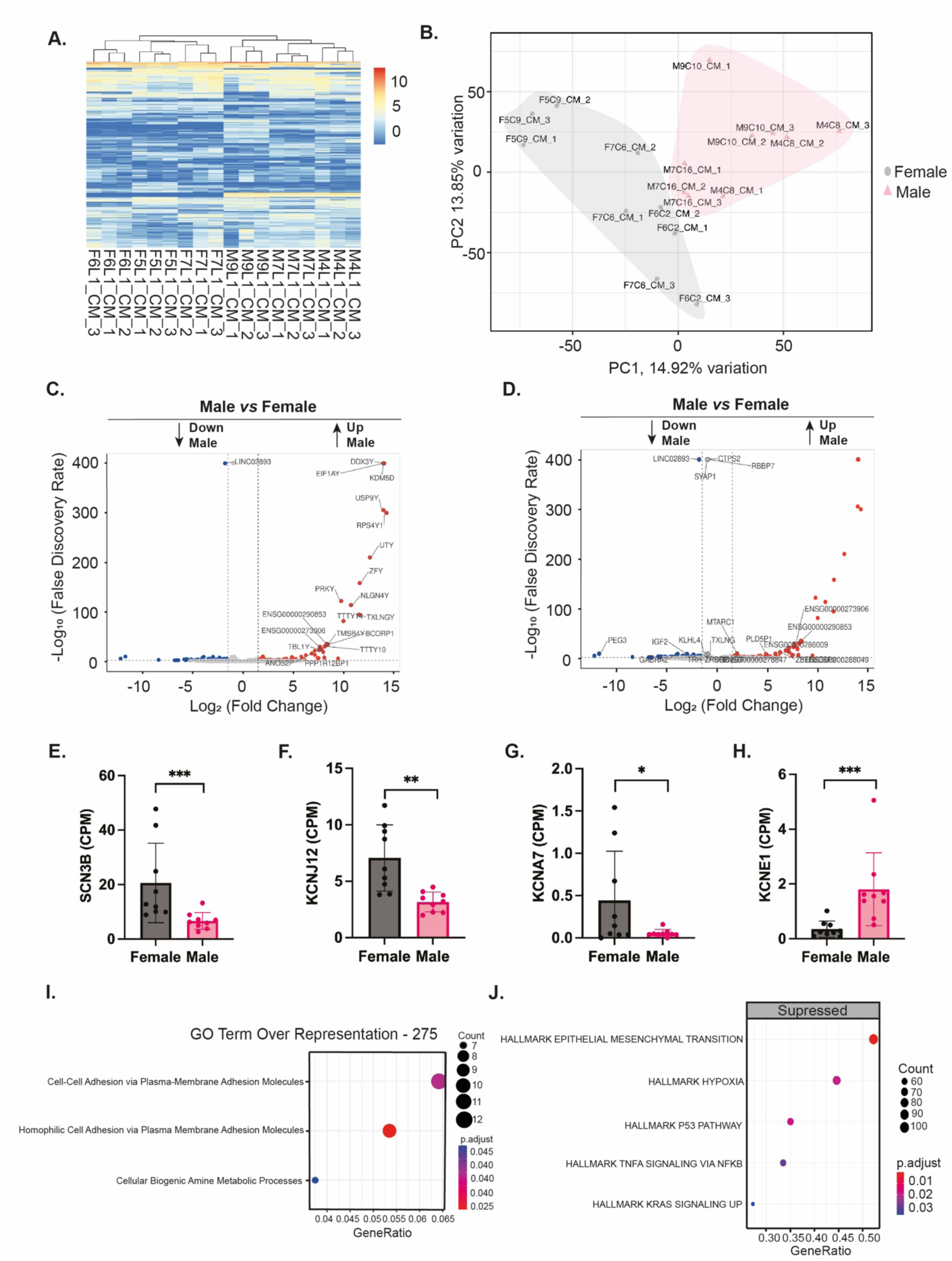
Bulk RNA sequencing reveals genetic differences between female and male hiPSC-CM. A) Heat map of hiPSC-CM gene expression from three independent experiments for all 6 hiPSC-lines. A dendrogram at the top of the heatmap shows separate hierarchical clustering of male and female hiPSC-CM based on gene expression patterns. B) Principal Component Analysis (PCA) plot showing separate clustering of male and female hiPSC-CM where each dot is one independent RNA sequencing replicate. C) Volcano plot representing differentially expressed genes (DEGs) between male and female hiPSC-CMs. D) Volcano plot analogous to (C) with the exclusion of known Y-linked genes for visualization of non-Y-linked genes on the volcano plot. Graphs showing the expression level in counts per million (CPM) for the ion channel genes E) sodium voltage-gated channel 3 B (SCN3B), F) potassium inwardly rectifying channel subfamily J member 12 (KCNJ12), G) potassium voltage-gated channel subfamily A member 7 (KCNA7), and H) potassium voltage-gated channel subfamily E regulatory subunit (KCNE1). I) Dot plot showing enriched gene ontology (GO) biological process (BP) pathways from an overrepresentation analysis of the DEGs between the male and female hiPSC-CM. The dot size indicates the number of genes linked to each enriched pathways, while the dot color reflects the statistical significance of the enrichment. F) Hallmark gene set enrichment dot plot highlighting gene sets that are suppressed in male hiPSC-CM compared to the female hiPSC-CM. The dot size signifies the number of genes associated with enrichment, and the dot color denotes the level of statistical significance. For (E-G) the bar represents the average ± STDEV and each dot is the normalized expression level in CPM for one independent experiment with n = 3 replicates per lines and three hiPSC-CM lines per sex. Statistical significance designations are based off an Edge Test of raw reads and the False Discovery Rate corrected p-value. Where *p < 0.05, **p < 0.01, ***p < 0.001, and ****p < 0.0001.

Principal Component Analysis (PCA) showed distinct clustering of the male and female-derived samples **(Figure 4B).** Notably, experimental replicates from the same cell line formed tighter clusters with each other, highlighting the batch-to-batch reproducibility of the data.

Differentially expressed genes (DEG) between the female and male hiPSC-CM were determined by grouping all the samples from each line together by sex. The female expression data was used as the reference for calculating the fold-change in gene expression between the male and female hiPSC-CM. There were 347 DEG with a false discovery rate (FDR) corrected p-value < 0.05.

These were plotted as a volcano plot with the genes upregulated in female hiPSC-CM and downregulated in male hiPSC-CM having a negative fold change; conversely, the genes with a positive fold change were upregulated in male hiPSC-CM and downregulated in female hiPSC-CM (**Figure 4C**). DEG were also evaluated with known Y-linked genes removed highlighting the top non-Y-linked DEG in the male and female-derived CMs (**Figure 4D**).

Due to differences in calcium transients, and previous reports of different potassium ion channel compositions in hiPSC-CM, the list of DEGs was probed in a biased manner for any calcium handling or ion channel genes (**Figure 4**). There were no calcium-handling genes identified, corroborating previous reports that the majority of calcium-handling differences are mediated by estrogen^34,35^. However, we found differential expression of four ion channels relevant to the cardiac action potential and pertinent to cardiac health and disease. Female hiPSC-CM showed increased expression of 1) sodium voltage-gated channel beta subunit 3 (SCN3B), 2) potassium inwardly rectifying subfamily J member 12 (KCNJ12), and 3) potassium voltage-gated subfamily member 7 (KCNA7) (**Figure 4E-G**). Differential expression of these ion channels in a healthy hormone-free system has, to our knowledge, never been reported. We also validate here an increase in potassium voltage-gated channel subfamily E regulatory subunit 1 (KCNE1) expression in male hiPSC-CM reported in one previous study (**Figure S4H**)^30^. These results indicate baseline differences in ion channel composition between male and female hiPSC-CM and serve as rationale for the inclusion of sex in pharmacological studies that target these channels and other channels the cooperate to generate the cardiac action potential. Next, DEG were examined in an unbiased manner using gene ontology (GO).

### Gene ontology pathways enrichment reveals sex differences in CM cell adhesion and amine metabolism

To better understand the differences in gene expression between male and female hiPSC-CMs, GO overrepresentation analysis was used to find out if any pathways were enriched between the female and male hiPSC-CM. Both “Cell-Cell Adhesion via Plasma-Membrane Adhesion Molecules” and “Homophilic Cell Adhesion via Plasma Membrane Adhesion Molecules” were enriched GO biologic process (BP) pathways, processes that are crucial for cell interaction and communication (**Figure 4I**). Additionally, the pathway “cellular biogenic amine metabolic process” was also enriched. Using a computer network plot (cnet) we visualized the connections between specific genes that were responsible for the GO term enrichment of both the cell adhesion pathways and the amine metabolism pathway (**Figure S4 and S5**). To determine which genes were up and down-regulated between the male and female hiPSC-CM, bar plots of the normalized counts per million (CPM) of all the genes in each pathway were plotted. We found that the hiPSC-CM had differential expression of multiple cadherins (CDH), cell adhesion molecules that serve to mechanically connect cells. Female hiPSC-CM had increased expression of CDH1 and CDH17 while male cells showed increased expression of CDH7 and CHD2 (**Figure S4B-M**). These results indicate that male and female hiPSC-CM potentially adhere to each other in using different proteins even in the absence of exogenous hormones. Thus, differences in cell-cell adhesion may be a potential mediator of non-hormone-mediated sex dimorphism in cardiac health and disease.

When looking at DEG for the amine metabolism pathway, a large and distinct increase in thyrotropin-releasing hormone (TRH) was revealed in all the replicates and lines for the female hiPSC-CM (**Figure S5B**). TRH is a cardiac hormone that has important functions in heart regulation and the responses to cardiac stress. It is induced after myocardial infarction and serves as a positive inotrope increasing cardiac contractility and output, but long-term activation can lead to a hypertrophic phenotype in the heart^42–45^. To further interpret the gene expression data and enhance the GO analysis, a hallmark gene set analysis was conducted with female-derived hiPSC-CMs as a reference.

### Hallmark gene sets related to hypoxia were suppressed in male hiPSC-CM

Hallmark gene sets are groups of genes that each represent a distinct biological condition, process, or pathway and tend to be active simultaneously^46^. In male hiPSC-CMs, there is a notable suppression in the activity of the hallmarks of “Epithelial Mesenchymal Transition,” “Hypoxia,” “TNFA Signaling via NFKβ,” “P53 Pathway,” and “KRAS Signaling Up” gene sets relative to their female counterparts (**Figure 4J)**. This reflects potential variations in stress response and common signaling pathways wherein female cells show some level of basal activation. When looking at heat maps of the hallmark genes that caused pathways enrichment in each set by hiPSC-line it is apparent that these pathways tend to be more activated in female hiPSC-CM than the male hiPSC-CM (**Figure S6**). Additionally, multiple regions of interest can be identified in each heat map where all the female hiPSC-CM have higher expression than all of the male hiPSC-CM. For example, the genes for PGK1, SLC2A3, COL5A1, PKP1, PDK3 and NDRG1 in the hypoxia heatmap, the genes GFPT2, GYPC, THEM176A and THEM176B in the P53 pathways heatmap, the genes RNF19B, ZBTB16, TPD52L1, H2AJ and KRT17 in the TNFA signaling heatmap as well as the genes MAP3K8, AREG, CCRL2, DUSP2 and RNF19B all show a consistent expression pattern in all six hiPSC-lines CM. This result helps visually describe the robust differential expression of genes in the hallmark pathways related to cell signaling and responses to stress. All of the information gleaned from this analysis reveals potential targets for non-hormone-mediated sex dimorphism in hiPSC-CM. In total, this study reveals functional and associated genetic differences in hiPSC-CM from males and females making the inclusion of sex as a variable when studying hiPSC-CM *in vitro* essential.

## Discussion

Here, we first established sex as an underreported and understudied variable in hiPSC-CM primary research literature. We then established a resource for studying sex differences in hiPSC-CM by reprogramming three female and three male hiPSC-lines from adult left ventricular adult fibroblasts. When differentiated into hiPSC-CM we discovered a slight but significant increase in the net calcium handling ability of female hiPSC-CM in comparison to male ones. Additionally, the female CMs had increased irregularity of spontaneous beat rate. We established that female and male hiPSC-CM also had different patterns of gene expression with over 300 DEG leading to distinct clustering. GO pathways analysis determined the most enriched pathways between the male and female hiPSC-CM were those related to cell-cell adhesion and amine metabolism. Gene set enrichment determined significant suppression of pathways related to hypoxia, p53, KRAS, and TNFA signaling in the female hiPSC-CM. Together, these results reveal baseline sex differences in hiPSC-CM at the functional and transcript level without the addition of exogenous hormones.

Male and female hiPSC-CM showed differentiation expression of multiple cardiac ion channel that have implications for health and disease. First, we bolstered the findings of a previous research study that demonstrated increased expression of the KCNE1 gene in male hiPSC-CM; making the female hiPSC-CM more susceptible to proarrhythmias such as Torsades de pointes^30^. This study only looked at two male and two female lines, with only one replicate per line making these initial results tenuous with statistical significance difficult to establish. Our study confirms this finding across three male and three female lines with three independent hiPSC-CM differentiation batches for every line. The primary function of KCNE1 in CM is regulating the cardiac action potential duration^47^. It serves as a major contributor to the QT interval, or the repolarization current of CM^47^. As mentioned above, KCNE1 is implicated in genetic and drug Long-QT syndrome and Torsades de Pointes which are arrythmias more common in women^48,49^. These results confirm sex is an important variable to consider in pharmacological studies on hiPSC-CM, especially for drugs that act as potassium channel blockers.

We also discovered that females have significantly higher expression of the sodium ion channel SCN3B, and the potassium channels KCNJ12 and KCNA7. SCN3B is a sodium ion channel and increases in SCN3B gene expression have been correlated to increases in sodium current in myocardial cells^50^. Conversely, knockouts and mutations of the SCN3B gene in mice show decreased cardiac sodium current density and abnormal electrophysiology^51,52^. Mutations have specifically been linked to the onset of atrial fibrillation, an arrhythmia more prevalent in men^7,51^. The lower expression of SCN3B in non-diseased male CM could be a mediator of this disparity which provides the scientific premise for future research. Brugada syndrome can cause sudden cardiac death, and is also more common in men; this disease tends to be masked in children with the risk of sudden cardiac death increasing substantially with age^7,53^. Recent studies using hiPSC-CM postulate that the fetal and post-natal expression of SCN3B masks the Brugada phenotype at early ages^53^. Our results, showing an increase in SCN3B in female hiPSC-CM, could explain some additional sex-specific masking of the Brugada phenotype in the female sex, leading to the decreased prevalence of sudden cardiac death in females with the Brugada genotype. Another key ion channel, we found to be present at higher levels in female hiPSC-CM, was KCNJ12. Expression of this gene is decreased in hearts with dilated cardiomyopathy in comparison to healthy controls, making decreases in KCNJ12 expression either a consequence of the disease or a potential mediator of it^54^. Interestingly, non-genetic dilated cardiomyopathy is a more common phenotype seen in men with heart failure^6^. Taken together these results indicate sex differences in ion channel expression in cardiac health, could be mediators of sex differences in cardiac disease.

Our investigation also highlights hiPSC-CM baseline sex differences in calcium handling, somewhat corroborating existing findings regarding sex differences in human and animal hearts. One study showed that female rat hearts expressed significantly more ryanodine calcium release channel (RyR), the L-type calcium channel IC (CANCA1 or Cav1.2), and the sodium-calcium exchange channel (NCX)^55^. Later studies attributed these differences to sex hormones. A study looking at rabbit hearts saw an increase in calcium handling spurred by increases in L-type calcium channels and sodium-calcium exchanges in the apex of female rabbit hearts^34^. When the cells were excised, the exogenous addition of estrogen led to a doubling in the calcium current in CM from the rabbit heart apex. This was later validated in primary human ventricle tissue where female apex tissue had a higher expression of L-type voltage-dependent channel Cav1.2 and the sodium-calcium exchanger NCX1^35^. This result was seen only in premenopausal women indicating estrogen might play a role. When male and female hiPSC-CM were conditioned with estrogen both the sodium and calcium current increased in the female CM but not the male^35^.

Our findings confirm that hormones are likely the cause for the differential expression of Cav1.2, RYR and NCX as none of those genes were differentially expressed in our dataset. However, our findings also suggest that female CM have a slight but significant increase in calcium handling even without the addition of exogenous hormones. The etiology for these changes could be linked to differential expression of other ion channels or a yet undiscovered mechanism.

Notably, we also observed significant differences in cell-cell adhesion. Cell adhesion processes are not just mechanical connections between cells but are also crucial for the transmission of signals between cells. These are especially crucial in highly mechanically active organs such as the heart^56^. In our data set, DEGs in these pathways revealed differences in female and male cadherin (CDH) expression. In our data set CDH1 and CDH17 were upregulated in female hiPSC-CM while CDH7 and CDH12 were upregulated in male hiPSC-CM. Although N-cadherin (CDH2) is more abundant in the heart, and plays a critical role in intercalated disk assembly, CDH1 can also be found in the heart^57,58^. In a genetic rat model of cardiac hypertrophy, the hypertrophic rats showed significantly higher levels of CDH1 compared to healthy rats and this expression was particularly concentrated at the CM intercalated discs^58^. When CDH1 was overexpressed in rat and rabbit myocytes it led to reduced cell size and the movement of β-catenin out the nucleus, thus inactivating WNT signaling^58^. This suggests that altered cell adhesion, through the expressions of cadherins, may play a role in cardiac hypertrophy. The basal differences in cadherin expression between male and female hiPSC-CM could be mediators of sex dimorphism in cardiac hypertrophy and merit future study^59^.

DEG from the Bulk RNA Sequencing Analysis was used to create a Volcano Plot, visually representing the gene expression variations between cardiomyocytes derived from males and females. As expected, some of the DEG were located on the Y chromosome, which is exclusive to males. The role of Y-linked genes in cardiac health and disease is significant. For instance, an expression profiling study of patients with new-onset heart failure due to idiopathic dilated cardiomyopathy identified a marked upregulation in USP9Y, DDX3Y, RPS4Y1, and EIF1AY, correlating with pronounced fold changes in male patients^60^. Additionally, research comparing control subjects to heart failure patients found that out of seven highly expressed gene IDs in the heart failure group, five were Y-linked genes: EIF1AY, RPS4Y1, USP9Y, KDM5D, and DDX3Y^61^. All of these Y-linked genes were significantly increased in the male hiPSC-CM of this study. The precise contributions of these Y-linked genes to cardiac health and disease remain under-explored, presenting another potential avenue for future research.

The separate bulk RNA sequencing clustering of the male and female hiPSC-CM is significant as other studies have determined that, even in large cohort studies encompassing hiPSCs from 40 donors, hiPSCs themselves cluster independently of sex and age^62^. This is significant to note as it shows that the inherent cardiac sex dimorphism of hiPSC-CM is great enough to be detected even with a cohort of six lines. Though some of the most significantly differentially expressed genes are Y-linked the sex differences are not necessarily seen in all cell types but are unique in this case to the hiPSC-CM themselves.

Of note, the hiPSC-CM of this study exhibit maturity levels of the fetal period^63^. Future studies could impose maturation conditions on the hiPSC-CM, such as metabolic maturation media, electrical stimulation or 3D engineered heart tissue culture^64–68^. A recent review eloquently laid the framework for the future of studying sex in more complex engineered tissues, often including multiple cell-types^69^. In addition to studying the phenotype after maturation conditions are imposed, mechanistic information can be gleaned as to how female and male CM mature. This is highly relevant as there is sex dysmorphism in the presence and frequency of various congenital heart defects^70^. Additionally, a recent study looking at single cell RNA sequencing data at different developmental time points showed that cardiomyocytes had the most DEG at multiple developmental time-points, and that this was largely mediated by sex-specific progesterone receptor differences driving maturation^36^. Another future consideration would be to add hormones. By not adding hormones, we were able to see baseline sex-differences in the hiPSC-CM that are not driven by hormones. Future studies might include the addition of hormones since, as described above, estrogen plays a key role in cardiac development, health and disease.

A challenge of working with human source material is line-to-line variability which limits our ability to detect subtle differences between sex. One approach to address this issue is to increase the number of male and female lines. There was also a recent study that generated isogenic hiPSC-lines with various sex chromosomes; XY, XX, XXY and X0. In the future, banks of lines such as these could be used to study sex differences without differences in non-sex chromosome genetic background^71^. This also brings up the fact that our study only looked at the dominant sex chromosome orientations, XX and XY, where there could be other basal differences in individuals with Klinefelter syndrome (XXY), Turner syndrome (X0) or other sex chromosome abnormalities. These other sex chromosome variants should be considered in future studies.

Along the lines of sex chromosomes, the hiPSC somatic cell reprogramming process is limiting in itself because of the high tendency for hiPSC to contain irregularities in X-chromosome inactivation^72,73^. This is limiting when studying sex differences using hiPSC-derived cells as genes that escape X-inactivation can be mechanistic mediators of cell-level sex dimorphism^74,75^. Last year it was discovered that reprogramming hiPSCs through a naïve state highly reduced abnormalities in hiPSC X-inactivation along with other epigenetic abnormalities acquired with reprogramming^76^. In the future adoption of this, or similar reprogramming methods, could be used when studying sex differences using hiPSCs.

Overall, our study demonstrates that sex-related differences are present in hiPSC-CM even without the addition of exogenous hormones. This was achieved using adult left ventricular fibroblasts with the most common Sendai virus reprogramming technology, in an attempt to limit epigenetic differences in cell sourcing and reprogramming. These findings highlight the importance of considering sex as a variable in hiPSC-CM and identify novel mediators of cardiac sex dimorphism that are not mediated by exogenous sex hormones. So that these outcomes and others related to CVD can be more easily studied, we provide an essential resource for the field in the form of a controlled cohort of male and female hiPSCs.

## Materials and Methods

### Literature Search

The goal of this literature search was to analyze publications from PubMed between 2010 and 2023 to assess whether sex as a biological variable has been considered in hiPCS-CM research. We began broadly, using PubMed to identify papers on hiPSC-CMs with the search term “(Human Induced Pluripotent Stem Cell derived Cardiomyocytes) OR hiPSC-CMs OR (human iPSC-CMs)”. We tracked the annual count of such papers from 2010 to 2023. Next, we refined our search to include “(sex OR male OR female)” to determine whether any of these terms were mentioned in the context of hiPSC-CMs. To hone in even further, we refined our search to include “(sex AND male AND female)”, to determine if any hiPSC-CM publication contained all of these terms. Recognizing the limitations of this approach, we conducted a more thorough search. We input “(Human Induced Pluripotent Stem Cell derived Cardiomyocytes) OR hiPSC-CMs OR (human iPSC-CMs)” into PubMed and reviewed the first 50 primary research articles listed for each year from 2010 through 2023. In each publication reviewed we determined whether the sex of the hiPSC-CM cell lines was reported and then whether the results were analyzed by sex.

### Adult left ventricular cardiac fibroblasts (aLVCF) isolation

All human tissue procurement and use was performed with local Institutional Review Board approval. Donor human hearts that were considered healthy based on available clinical information, but still went unmatched for transplant were obtained from the University of Wisconsin Organ Procurement Organization, Madison, WI. Donor hearts associated with underlying coronary artery disease, impaired left ventricular function (ejection fraction <50%), known bacteremia, viral hepatitis or HIV, or anticipated cold ischemic time of >6 hours were excluded. The aLVCF isolation was performed as previously described^77–80^. Briefly, hearts were flushed with ice-cold cardioplegia solution, excised and immediately submerged in ice-cold cardioplegia solution, and transported on ice. 25-30 g of left ventricle was excised, minced, distributed equally into 4 gentleMACS C-tubes, and homogenized with a gentleMACS tissue dissociator (Miltenyi Biotec, Bergisch Gladbach, Germany). 10 ml of digestion media containing DMEM (Corning, Corning, NY) and 2.5 mg Liberase TM (Roche, Basel, Switzerland) were added to each tube and incubated at 37 °C for 30 min with constant agitation. Heart samples were sieved through a 200 μm filter and then centrifuged at 1000×g for 20 min. The cell pellet was suspended in 20 ml complete media (MCDB 131 with 10% FBS (R&D Systems, Lot E20074), 1 ng/ml basic fibroblast growth factor (bFGF), 5 μg/ml insulin, 10 μg/ml ciprofloxacin and 2.5 mg/ml amphotericin B) and plated into two T75 flasks. Cells were permitted to attach for 2 hours, which is a time that offered an optimal CF yield with high purity during earlier protocol optimization studies. Non-adherent cells were removed by washing in 1X phosphate-buffered saline (PBS). Finally, complete media was added, and the cells were cultured at 37 °C, 5% CO2, and 100% humidity, and passaged once they were ∼90% confluent.

### LVCF reprograming into human-induced Pluripotent Stem Cells (hiPSCs)

The aLVCF lines were obtained via MTA (MSN251847) using an IRB-governed protocol at the University of Wisconsin-Madison. For this study, six hiPSC lines were used, all originating from left ventricular cardiac fibroblasts. Three lines were from female donors (F7, F6, and F5-LVCFs) and three lines from male donors (M4, M7, and M9-LVCF) between the ages 21 and 34 with no cardiac abnormalities (**Table S1**). The LVCFs were reprogrammed using the Cyto-tune 2.0 (Thermo A16517) Sendai Reprogramming Kit according to the suppliers’ directions. Briefly, low passage LVCFs, between passages 1 and 3, were plated at a density of ∼7000 cell/cm^2^ in LVCF-media (MCDB131 Low Glucose, 10% FBS, 5 μg/mL recombinant human insulin and 1 ng/mL recombinant human basic fibroblast growth factor). The following day Cyto-tune 2.0 Sendai virus was added to the LVCF-media and removed 24-hours later. The cells were maintained in LVCF-media for 6 more days. On day 7, the cells were singularized using Accutase and replated at a 1:6 ratio onto a vitronectin (Thermo A31804) coated plate. To coat plates with vitronectin, the vitronectin was diluted to a 10μg/mL concertation in DPBS minus calcium and magnesium and left at 4°C overnight or at least 1-hour at room temperature. The day following replating, the media was changed to TesR-E8 (STEMCELL 05990), and the cells were maintained in TesR-E8 for the next 2-3 weeks while colonies began to emerge.

Independent colonies that looked pluripotent were manually passaged using a microscope and Pasteur pipette tip in the biosafety hood. Approximately 12-18 colonies were selected for each line and each colony was manually passaged 5-times to aid exogenous Sendai vector elimination. After 5-manual passages colonies that were maintaining a pluripotent colony phenotype were selected and bulk passaged using Versene (Thermo A423101) onto vitronectin-coated plates.

Between passage 8-20 6 colonies were tested via q-RT-PCR for Sendai vector using the primer pair recommended in the Cyto-tune manual (**Table S2**). A line was considered Sendai vector depleted when no SEV product was detected for two passages via qPCR for 40-cycles. Three clones that were Sendai vector depleted were expanded and banked using mFresR (STEMCELL 05855). One Sendai vector-depleted clone from each line was chosen to move forward with for comprehensive pluripotency assessment, karyotyping, and cardiomyocyte differentiations (F7C6, F6C2, F5C9, M4C8, M9C10, and M7C16-hiPSCs). For ease of use the lines later remained as the donor identifier (e.g. F5) followed by L1, L2, or L3 for lines 1, 2, and 3 as shown in **Table S1**.

### Pluripotency assessment

The pluripotent status of the hiPSC-lines was assessed via q-RT-PCR, flow cytometry, and immunocytochemistry (ICC). For q-RT-PCR the cells were lysed, and RNA was purified using the PureLink^TM^ RNA Mini Kit (Thermo 12183025). The purified RNA was converted to cDNA using SuperScript^TM^ IV VILO^TM^ Master Mix. The qPCR was completed using forward and reverse primers for OCT4, SOX2, and NANOG with SYBR Green PCR Master Mix (Thermo 4367659). The primer sets are shown in **Table S2**. The βCt was calculated in comparison to the reference gene GAPDH and was compared to aLVCF as a negative control and a previously characterized hiPSC-line, CCND2-hiPSCs as a positive control (Insert CCND2-hiPSC Reference). To confirm the presence of pluripotency markers at a protein level flow cytometry was for OCT4 and SSEA4 with the detailed methods outlined below. Additionally, four pluripotency markers, OCT4, SSEA4, SOX2, and TRA-1-60 using ICC following the suppliers’ directions from the Pluripotent Stem Cell 4-Marker ICC Kit (**Table S3**)

### hiPSC-lines Karyotyping

Adherent cells were harvested with colcemid arrest (Irvine Scientific), treated with 0.75 M KCl hypotonic solution, and fixed with 3:1 methanol:acetic acid. The resulting cells were spread onto glass slides according to standard cytogenetic protocols, aged in a 90-degree oven for 1.5 hrs and stained with Wright’s/Giemsa stain. Twenty G-banded metaphases were evaluated using an Olympus BX51 microscope outfitted with 10x and 100x objectives and karyotyped using Applied Spectral software. Olympus BX51 with 60x and 100X oil objective lenses.

Interferometer-based CCD cooled camera. Software for G-bands and SKY: Applied Spectral Imaging (ASI) BandView.

### Cell maintenance of hiPSCs

The hiPSCs were cultured in mTesR^TM^1 (STEMCELL 85850) on 8.68 ug/cm^2^ (or 0.5 mg per 6 well-plate) Cultrex reduced growth factor (RGF) (R&D Systems 3433-005-01) coated 6-well plates. The media was changed into 2mL of fresh mTesR every day. Upon reaching 70-80% confluency, hiPSCs were passaged using ReLesSR^TM^ (STEMCELL 05872). The exhausted media was aspirated, and the hiPSCs were washed with 1 mL of DPBS (-) calcium (-) magnesium to remove residual medium and contaminants. Then, 1 mL of room temperature ReLesSR™ was added to each well, followed by a 45-second incubation. After aspirating the ReLesSR™, the plate was placed in a 37°C incubator for 7 minutes. After the incubation, 1 mL of warm mTeSR™1 was added to each well, and the hiPSC colonies were detached from the bottom by gently tapping the sides of the plate. The cell suspension was diluted at a 1:6 ratio and transferred into a new Cultrex-RGF coated 6-well plate at 2mL cell suspension per well.

### Differentiation and purification of hiPSC-derived Cardiomyocytes (hiPSC-CMs)

Upon reaching 70-80% confluency, hiPSCs were dissociated using Accutase solution (Millipore Sigma A6964). The exhausted media was aspirated, and cells were washed with 1 mL of DPBS (-) calcium (-) magnesium. Then, 1 mL of room temperature Accutase solution was added to each well and incubated at 37°C for 8 minutes. The hiPSCs were then singularized by pipetting up and down with a P1000 micropipette and quenched using ½ mL of mTesR. The resulting cell suspension was transferred to a conical tube and counted on a hemocytometer using a 1:1 dilution of 0.4% trypan blue. The hiPSCs were then seeded onto a 12-well plate coated with 8.68 ug/cm^2^ Cultrex-RGF at a density of 0.5×10^6^ cells per well in 1 mL of mTesR with 5uM Rock Inhibitor. The exhausted media was replaced every day with 2 mL of fresh mTesR until the cells reached 97-100% confluency (2-4 days). Upon reaching confluency (defined as day 0 of differentiation), the media was replaced with RPMI/B27(-insulin) (Fisher A1895601) containing varied CHIR99021 (Sigma SML1046) concentrations based on the hiPSC line (**Table S1**). On day 2, exactly 48 hours later, the media was replaced with RPMI/B27(-ins) containing 7.5 uM of IWP2 (Tocris 686770-61-6), followed by another media change with RPMI/B27(-ins) on day 4. The media was then replaced every two days until day 13 of differentiation, using fresh RPMI/B27(+ins) (Thermo 17504001).

On day 13, the cardiomyocytes were replated onto 8.68 g/cm^2 Cultrex-RGF coated 12-well plates at a 1:2 ratio to facilitate lactate purification. Each well was washed with 1/2 mL of DPBS (-) calcium (-) magnesium. Subsequently, 1 mL of 37°C 0.25% trypsin-EDTA solution was added to each well and incubated at 37°C for 20 minutes. After incubation, cardiomyocytes were singularized by pipetting with a P1000 micropipette, then quenched with 2 mL of RPMI/B27(+ins) with 20% FBS. The resulting cell suspension was transferred to a conical tube, centrifuged at 200g for 5 minutes, and the supernatant was aspirated. Cells were resuspended in RPMI/B27(+ins) containing 10 uM Rock Inhibitor and seeded at 1mL cell suspension per well. On day 14, the replated cells were given 1 mL of RPMI/B27(+ins). Cardiomyocytes were then purified using lactate purification media composed of DMEM no glucose with 4 mM lactate. On day 15, the media was replaced with 1 mL lactate medium. On day 17, the cells were washed with 1/2 mL of DPBS (-) calcium (-) magnesium, and the media was replaced with 1 mL of lactate medium. On day 19, the cells were washed with 1/2 mL of DPBS (-) calcium (-) magnesium and switched into RPMI/B27 (+ins). The media was changed every three days with fresh RPMI/B27(+ins) until the cardiomyocytes were lifted for flow cytometry and RNA lysis on day 40, or for Calcium Imaging on day 60.

### Flow Cytometry

Flow cytometry was used to assess hiPSCs pluripotency and hiPSC-derived cardiomyocyte purity after lactate purification. For cell fixation and permeabilization, the hiPSCs were singularized using Accutase (the same method as used above). After incubation, the cells were quenched with 1/2 mL of mTeSR per well, and the resulting cell suspension was centrifuged for 5 minutes at 200g. The supernatant was removed, and the cells were fixed in 1% paraformaldehyde for 20 minutes and centrifuged again for 5 minutes at 200g. The PFA was removed, and the cells were resuspended in 90% v/v methanol and stored at -20°C for up to 6 weeks. For cardiomyocytes, on day 40, each well was washed with 1/2 mL of DPBS (-) calcium (-) magnesium and incubated with 1/2 mL of 37°C 0.25% trypsin-EDTA for 20 minutes at 37°C. After incubation, the cardiomyocytes were singularized with a P1000 micropipette and quenched with 1 mL of RPMI/B27(+ins) with 20% FBS. The cells were transferred into a conical tube and centrifuged for 5 minutes at 200g. The supernatant was removed, and the cardiomyocytes were fixed in 1% paraformaldehyde (PFA) for 20 minutes and centrifuged for 5 minutes at 200g. The PFA was removed, and the cells were then resuspended and stored in 90% v/v methanol at -20°C for up to 6 weeks. For flow cytometry stains the fixed cells were washed out of the methanol with 2 mL of flow buffer (PBS with 0.1% Triton-X and 0.5% w/v BSA) twice. The cell pellet was then resuspended in 100uL of primary antibody at the dilution indicated in **Table S3**. The fixed cells were incubated in primary antibody for 1 hour at room temperature and then washed twice with 2 mL of flow buffer. If the primary antibody was conjugated to a fluorophore the control was an isotype conjugated to the same fluorophore and the cells were resuspended in 300uL of flow buffer and 10,000 events were acquired on a BD Accuri C6 Flow Analyzer. For stains with unconjugated primary antibodies, the cell pellet was resuspended in 100uL of secondary antibody and incubated at room temperature for 30 minutes. The fixed cells were washed twice with 2mL of flow buffer, resuspended in 300uL of flow buffer, and then 10,000 events were acquired on a BD Accuri C6 Flow Analyzer. The negative control for the cardiac troponin T stains were undifferentiated hiPSCs.

### Calcium transients of Cardiomyocytes

On day 60, cardiomyocytes from each cell line were incubated with 5uM Rhod-2AM calcium-sensitive dye in LASR medium for 30 minutes at 37°C. Following the incubation period, the dye was removed and replaced with Tyrode’s Salt Solution containing 1 g/L sodium bicarbonate (TSS, Sigma Aldrich T2145). The cells were allowed to equilibrate in TSS for 30 minutes at 37°C and were subsequently subjected to imaging on the TxRed channel using a 30ms exposure and a gain of 1.0 at 10X magnification on a fluorescent microscope (Leica Microsystems). For each well, three spontaneous and three-paced 20-second videos were captured, with n=3 wells per condition for each replicate. For the videos taken with electrical stimulation, the cells were paced using a platinum electrode at 1 Hz with a voltage of 7 V and a pulse duration of 0.02 ms. The calcium transient videos acquired were processed using ImageJ. Three regions of interest within each video were selected, and their respective Z-axis profile was saved as .txt files. The text files were analyzed using a MATLAB code to determine the inter-spike interval, time to peak, maximum amplitude, as well as the average upstroke and downstroke velocities.

### Bulk RNA Sequencing

On day 40, cardiomyocytes were lifted in the same manner as for flow (described above). The cell pellet was quickly resuspended in a solution of 700μL PureLink Lysis buffer containing 7μL Beta-mercaptoethanol. The cell lysis was quickly transferred to -80°C for storage. The RNA purification was done using the PureLink^TM^ RNA Mini Kit (Thermo 12183025). The DNA was removed using the on-column Ambion PureLink DNase Set (Thermo 12185010). Total eukaryotic RNA isolates are quantified using a fluorimetric RiboGreen assay. Total RNA integrity is assessed using capillary electrophoresis (e.g. Agilent BioAnalyzer 2100), generating an RNA Integrity Number (RIN). For samples to pass the initial QC step, they need to quantify higher than 500 ng and have a RIN of 8 or greater. Total RNA samples are then converted to Illumina sequencing libraries. Total RNA samples are converted to Illumina sequencing libraries using Illumina’s TruSeq RNA Sample Preparation Kit (Cat. # RS-122-2001 or RS-122-2002) or Stranded mRNA Sample Preparation Kit (Cat. # RS-122-2101). (Please see www.illumina.com for a detailed list of kit contents and methods). In summary, the mRNA from a normalized input mass of total RNA is isolated using oligo-dT coated magnetic beads, fragmented, and then reverse transcribed into cDNA. The cDNA is blunt-ended, A-tailed, and indexed by ligating molecularly barcoded adaptors. Libraries are amplified using 15 cycles of PCR. The final library size distribution is validated using capillary electrophoresis and quantified using fluorimetry (PicoGreen) and via Q-PCR. Indexed libraries are then normalized, pooled, and size-selected to 320 bp (tight) using the PippinHT instrument. Pooled libraries are denatured and diluted to the appropriate clustering concentration. The libraries are then loaded onto the NovaSeq paired end flow cell and clustering occurs on board the instrument. Once clustering is complete, the sequencing reaction immediately begins using Illumina’s 2-color SBS chemistry. Upon completion of read 1, a 7 base pair index read is performed in the case of single indexed libraries.

If dual indexing was used during library preparation, 2 separate 8 or 10-base pair index reads are performed. Finally, the clustered library fragments are re-synthesized in the reverse direction thus producing the template for paired end read 2.

### Bulk RNA Sequencing Analysis

2 x 50bp FastQ paired-end reads for 6 samples (n=62.4 Million average per sample) were trimmed using Trimmomatic (v 0.33) enabled with the optional “-q” option; 3bp sliding-window trimming from 3’ end requiring minimum Q30. Quality control on raw sequence data for each sample were performed with FastQC. Read mapping was performed via Hisat2 (v2.1.0) using the Human genome (GRCh38) as a reference. Gene quantification was done via Feature Counts for raw read counts. Differentially expressed genes were identified using the edgeR (negative binomial) feature in CLCGWB (Qiagen, Redwood City, CA) using raw read counts. We filtered the generated list based on a minimum 2x Absolute Fold Change and FDR corrected p < 0.05. Differentially Expressed Genes (DEGs) were sorted based on log2(fold change) and -log10(false discovery rate, FDR). These values were then visualized in a Volcano Plot using the VolcaNoseR tool. Known Y-linked genes were filtered using the National Center for Biotechnology Information’s Gene database. Over-representation Analysis (ORA) and Gene Set Enrichment Analysis (GSEA) were performed on the differential gene expression results table(s) using the ClusterProfiler_wrapper v2.1 module at the Minnesota Supercomputer Institute. This script is a wrapper for the clusterProfiler R package v4.7.1.003 for R v4.2.2^81,82^. Over-representation analysis was performed on 275 DEGs genes after applying an FDR cutoff of 0.05, a log2(fold change) cutoff of 1, and a top number of genes cutoff of 400. The gene SYMBOLs were converted into ENTREZ IDs using the bitr() function for compatibility with the enrichGO (ontology=biological process, pvalueCutoff =0.05, qvalueCutoff = 0.2 ) and enrichKEGG (pvalueCutoff =0.05, qvalueCutoff = 0.2) clusterProfiler functions, and the enrichPathway (pvalueCutoff =0.05, qvalueCutoff =0.2) ReactomePA R package function^83^. The human genome-wide annotation database, ‘org.Hs.eg.db’ v3.16.0 R package, was used for gene conversions and the enrichment functions. The universe or background gene list that was provided to the enrichment functions included all tested genes in the differential gene expression results table. All of the tested genes were ranked by their log2 fold change values, and GSEA was performed on the ranked list using hallmark database pathways (qvalue_threshold = 0.05) from the msigdbr R v7.5.1 package^84^. Genes permutations were performed for each pathway 10,000 times to calculate the normalized enrichment score (NES). Dotplots were generated with the ggplot2 R package.

## Acknowledgments

Marissa Macchietto from the Minnesota Supercomputing Institute for assistance in the bulk RNA sequencing analysis. The cytogenetic analyses were performed in the Cytogenomics Shared Resource at the University of Minnesota with support from the comprehensive Masonic Cancer Center NIH Grant #P30 CA077598. Three of the aLVCF were banked and provided by the Palecek lab and Marth Floy at the University of Wisconsin.

## Notes

### Competing Interest Statement

The authors have declared no competing interest.

